# In-cell detection of conformational sub-states of a GPCR quaternary structure: Modulation of sub-state probability by cognate ligand binding

**DOI:** 10.1101/2020.06.29.178137

**Authors:** Joel Paprocki, Gabriel Biener, Michael Stoneman, Valerica Raicu

## Abstract

While the notion that G protein-coupled receptors (GPCRs) associate into homo- and hetero-oligomers has gained more recognition in recent years, a lack of consensus remains among researchers regarding the functional relevance of GPCR oligomerization. A relatively recent technique, Förster resonance energy transfer (FRET) spectrometry, allows for the determination of the oligomeric (or quaternary) structure of proteins in living cells via analysis of efficiency distributions of energy transferred from optically excited fluorescent tags acting as donors of energy to fluorescent tags acting as acceptors of energy and residing within the same oligomer. In this study, we significantly improved the resolution of the FRET-spectrometry approach to detect small differences between the interprotomeric distances among GPCR oligomers with subtle differences in quaternary structures. We then used this approach to study the conformational substates of oligomers of sterile 2 α-factor receptor (Ste2), a class D GPCR found in the yeast *Saccharomyces cerevisiae* of mating type **a**. Ste2 has previously been shown to form tetrameric oligomers at relatively low expression levels (between 11 and 140 molecules/μm^2^) in the absence of its cognate ligand, the α-factor pheromone. The significantly improved FRET spectrometry technique allowed us to detect multiple distinct quaternary conformational substates of Ste2 oligomers, and to assess how the α-factor ligand altered the proportion of such substates. The ability to determine quaternary structure substates of GPCRs provides exquisite means to elucidate functional relevance of GPCR oligomerization.

## 1. INTRODUCTION

G protein-coupled receptors (GPCRs) are a family of membrane-bound proteins which recognize and respond to a gamut of extracellular stimuli, thereby modulating the dissociation of the trimeric G protein from the cytoplasmic side of the receptor and inducing a down-stream signaling event^1–2^. GPCRs have traditionally been divided into six different classes according to their amino acid sequence and functional similarity: Class A-rhodopsin-like receptors, Class B-secretin family, Class C-metabotropic glutamate receptors, Class D-fungal mating pheromone receptors, Class E-cAMP receptors, and Class F-frizzled (FZD) and smoothened (SMO) receptors^3^.

Mounting evidence generated using a plethora of experiments, both in vitro and in living cells, indicates that GPCRs may form homo- or hetero-oligomeric complexes^4–11^. However, the prevalence and functional significance of such oligomeric complexes remains a largely open question. For some GPCRs, such as those belonging to the Class C-metabotropic glutamate subfamily, homo or hetero-oligomerization is absolutely essential for activation of the receptor^12^. By contrast, a number of other GPCRs, e.g., the β_2_-adrenoceptor, have been shown to retain functionality in monomeric form^13–15^, although they are also capable of forming dimers and/or higher order oligomers^16^. It would appear from these varied results that the degree of oligomerization, as it relates to function, may be class-specific, if not receptor-specific, although this remains quite unclear. In this regard, reliable methods that can report on the quaternary organization (i.e., oligomer geometry and interprotomeric distances) of GPCRs and specify how these structures are altered in response to activation by ligand binding are needed for a better understanding of the physiological importance of GPCR oligomerization.

Fluorescence-based methods, particularly those which rely on Fӧrster Resonance Energy Transfer (FRET), have been very effective in quantifying the interactions of membrane receptors within living cells. In FRET studies, the membrane proteins of interest are tagged with two fluorescent labels: a “donor” (D) and an “acceptor” (A). If the two fluorescent molecules reside within 10 nm of one another, a radiationless transfer of energy can occur from an optically excited D molecule to an unexcited A molecule^17^. The efficiency of the radiationless energy transfer is strongly dependent on the distance between the fluorophores, and therefore quantification of the FRET efficiency occurring between membrane protein labels reveals information about the relative proximity, and thereby interactions, of the receptors themselves^5,18–22^. If prior knowledge exists regarding the oligomeric size (i.e., number of protomers) and geometry (i.e., relative distances between protomers), then average-based FRET approaches allow one to determine relative proportions of monomers and different sized oligomers^23–26^.

A more recent addition to the family of FRET methods, FRET spectrometry^27–28^, can provide the size and geometrical parameters of the underlying quaternary structure of the receptor of interest. In FRET spectrometry, pixel-level values of FRET efficiency are assembled into distributions, or FRET spectrograms, from which the predominant (i.e., most frequently occurring) FRET efficiency values are extracted. A collection of the dominant FRET histogram peaks is then assembled into another histogram, termed a *meta-histogram*, to further separate the information originating from mixtures of oligomers with different sizes and geometries. To determine the size and shape of oligomers from meta-histograms, an iterative fitting process is performed which requires rigorous tests of how well various oligomer models simulate the measured meta-histogram^27,29^.

A number of methods, both experimental (e.g., fluorescence spectroscopy and crystallography) and computer-based (i.e., molecular dynamics simulations), which are well suited to study the tertiary structure of membrane receptors have shown that individual GPCRs are not rigid structures, but dynamic in nature, switching between multiple conformational substates^30–33^. It is therefore reasonable to expect that these fluctuations in the tertiary structure would lead to various quaternary structure substates as well. However, previous FRET spectrometry studies have been limited to determining only an average or most probable quaternary structure^27,29^

In the work described herein, we have significantly improved the resolution of FRET spectrometry so that it can resolve even slight alterations in the distance between protomers within the quaternary structure (i.e., less than 1 Å). The necessary improvement in resolution and sensitivity was accomplished, firstly, by isolating membrane-only regions of interest in the acquired images, and further dividing these regions into smaller segments. In this way, we capture fluctuations in quaternary structure between regions of the membrane which were previously smoothed out in the process of data analysis. Secondly, we refined our previous method of estimating the receptor concentrations (using two different excitation wavelengths)^27,34^ which allowed us to select subsets of the FRET spectrograms from cells expressing receptors in a narrow concentration range, to further sample the fluctuations of the receptor quaternary structure. This data acquisition strategy, combined with other methodological improvements and implementation of a noise-filtering algorithm, enabled a new picture to emerge of the quaternary structure versatility, which has never been observed in living cells. Specifically, we detected the presence of several quaternary structure conformations (or substates) in a typical class D GPCR, the sterile 2 α-factor pheromone receptor (Ste2), in the presence and absence of its cognate ligand^27,35–40^.

Ste2 is found in the yeast *Saccharomyces cerevisiae*^41–44^ of the mating type **a**. The receptor binds the α-factor pheromone, which is secreted from cells of mating type **α**, and subsequently initiates the signaling response, leading to the mating of haploid **a** and **α** cells^45–47^. Previous publications revealed that the Ste2 receptor forms complexes as large as tetramers at low concentrations^6,48–49^, with some octamers forming at higher concentrations^27^. While Ste2 is well characterized as a prototypical model for extracellular sensing, previous studies mainly focused on determination of the quaternary structure of Ste2 in the absence of ligand and lacked the necessary resolution to detect changes in the conformation^27^.

In this study, we were able to resolve four different quaternary structure conformations (or substates) assumed by Ste2, which were characterized by different distances between the protomers within the oligomer. We attribute these findings to the fact that the individual protomers comprising the oligomer can exist in multiple semi-stable, low-energy-state conformations, which is known from studies focusing on tertiary structures of other GPCRs^2,50–54^. Another significant finding was that, upon addition of ligand, the relative abundance of the quaternary conformations was shifted from the Ste2 structure primarily populating substates characterized by the smallest interprotomeric distances to substates with larger distances. Therefore, we were able to successfully associate the activity level of the receptor with a highly resolved quaternary structure conformation. These studies may be expanded to include other GPCRs (or any other membrane receptor) which may be exposed to different natural and artificial ligands, and thus significantly aiding in the search for the physiological relevance of GPCR oligomerization.

## 2. MATERIALS AND METHODS

### 2.1 Sample Preparation

Baker’s yeast cells (*Saccharomyces cerevisiae*) were engineered to express the sterile 2 α-factor pheromone receptor (Ste2) fused to one of two different fluorescent tags, i.e. GFP_2_^55^ or YFP^56–57^ at position 304 in the Ste2 amino acid sequence as previously described in Ref.^27^. Yeast cells transformed with one or both of the plasmids were grown at 30 °C on an agar-based plasmid selective synthetic complete medium lacking uracil and/or tryptophan.

Cell imaging was performed using 35-mm glass-bottom dishes (P35G-0.170-14-C, MATTEK Corporation, Ashland, MA). Prior to cell addition, the dishes were coated with concanavalin A (Sigma Aldrich, St. Louis, MO) to immobilize cells on the glass. To coat the dishes, 100 μL of a 0.5 mg/mL solution of concanavalin A (in deionized water) was placed on the coverslip of each dish. The dishes were then covered and incubated for 30 min at room temperature to allow deposition of the concanavalin A to occur. After 30 min, any remaining solution was removed, and the dishes were allowed to dry for a 24 h period prior to addition of cells.

Cells were scraped from the agar-based selective medium and suspended in 1 mL of 100 mM KCL buffer (pH 7.0). 200 μL of cell suspension was then pipetted onto the coverslip region of a concanavalin A coated glass-bottom dish and allowed to incubate for a period of 10 min. After the 10 minute incubation period, the dish was washed three times with 100 mM KCl buffer in order to remove any unbound cells, leaving a single layer of adherent cells for imaging^58^. For experiments where ligand effects were probed, an additional cell preparation step was introduced, as follows: A purified α-factor pheromone suspension (Y1001, Zymo Research, Irvine, CA) was diluted to a working concentration of 10 μM in 100 mM KCl to achieve and maintain saturation binding of ligand to receptors present within sample cell membranes^59–60^. Cells were suspended in 200 μL of the α-factor solution and allowed to incubate at room temperature for 5 min prior to plating. The incubated cell suspension was then added to a coated dish and incubated for another10 minutes to allow for the cells to adhere. After the incubation period, the dishes with adhered cells which had been exposed to ligand prior to plating were washed 3 times with the 10 μM ligand/buffer solution. After washing, the cell coated dishes were taken for imaging on a two-photon optical micro-spectroscope developed as described in the following section.

### 2.2 Two-photon fluorescence micro-spectroscopy

Fluorescence images were acquired using a spectrally resolved two-photon optical micro-spectroscope consisting of a tunable femtosecond laser (MaiTai™, Spectra Physics, Santa Clara, CA), an inverted microscope (Nikon Eclipse Ti™, Nikon Instruments Inc., Melville, NY) equipped with an infinity-corrected, plan apochromat, oil immersion objective (100×, NA=1.45; Nikon Instruments Inc.), and an OptiMiS scanning/detection head (Aurora Spectral Technologies, Grafton, WI), as described previously^27,61^. The samples were scanned using a line-shaped excitation beam with a power of 0.2 mW/voxel and an integration time of 35 ms per pixel. Each field of view was scanned at two different excitation wavelengths, first at 930 nm and then 800 nm, for the purpose of obtaining the concentration of both acceptors and donors (see Section 2.4 below). The total time needed to complete both excitation scans, including laser wavelength tuning time between scans, was ~60 s. The output of each excitation scan resulted in a set of micro-spectroscopic images which typically contained 200 different wavelength channels of ~1 nm bandwidth; the size of the image for each emission wavelength channel was 440 × 300 pixels^27^. For one particular experiment which focused on samples with particularly low receptor concentrations, the wavelength channel bandwidth was increased to ~2 nm in order to increase the level of fluorescence signal relative to the readout noise of the detector. Since molecular diffusion could potentially change the distribution of receptors and their oligomeric sizes for a given pixel during the relatively long time elapsed between each scan, both the donor and acceptor concentrations were computed as an average over a given region of interest rather than at pixel level.

### 2.3 Calculation of apparent FRET efficiency and assembly of meta-histograms

The composite emission spectrum from each pixel in the micro-spectroscopic images of cells co-expressing Ste2-GFP_2_ and Ste2-YFP was deconvoluted into donor and acceptor components using a least-squares fitting algorithm along with separately determined elementary donor and acceptor emission spectra, as described elsewhere^62^. The elementary fluorescence spectrum of the donor (green fluorescent protein, GFP_2_)^55^ and the acceptor (yellow fluorescent protein, YFP)^56–57^ were determined by acquiring micro-spectroscopic images of cells expressing either Ste2-GFP_2_ or Ste2-YFP. Applying the unmixing procedure to each image pixel resulted in 2D maps of donor intensity in the presence of the acceptor, *k^DA^*, and acceptor intensity in the presence of the donor, *k*^*AD*62^. The *k^DA^* and *k^AD^* values are two coefficients multiplying each elementary spectrum composing the theoretical function used to fit the measured spectrum. The *k^DA^* and *k^AD^* values were multiplied with the area underneath their respective elementary spectrum (i.e., their spectral integrals) to obtain the total donor fluorescence emission in the presence of the acceptor, *F^DA^*, and total acceptor emission in the presence of the donor, *F^AD^* at each pixel within an image^6^. Spectral integrals for the donor (*w^D^* = 43.59) and acceptor (*w^A^* = 42.92) were found from averages of typical elementary spectra obtained over multiple experimental days. A single spectral integral was used across multiple experiments in order to be able to compare receptor concentrations across multiple experimental days.

The apparent FRET efficiency (*E_app_*) for a mixture of donors and acceptors, some of which may be incorporated within oligomeric complexes, is defined as:

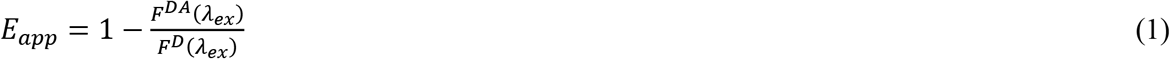

where *F^D^*(*λ_ex_*) is the fluorescence emission of the donor in the absence of the acceptor and may be expressed as follows^6,27^:

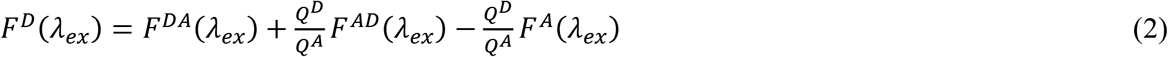

where *Q^D^* = 0.55 is the quantum yield of the donor^55^, *Q^A^* = 0.61 is the quantum yield of the acceptor^63^, and *F^A^*(*λ_ex_*) is the fluorescence emission of the acceptor in the absence of the donor^27^. Eq. (1) can be further simplified if an excitation wavelength is chosen for which the acceptor is negligibly excited (i.e., *λ_ex_* = 800 nm for YFP). When direct excitation of the acceptor is negligible, and hence *F^A^*(*λ_ex_*) ≈ 0, the apparent FRET efficiency can be found from the measured values of the donor and acceptor fluorescence, as follows^6^:

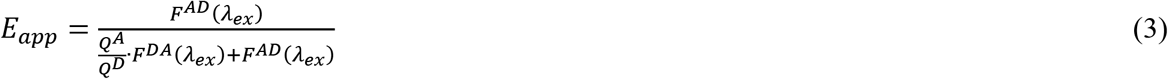

A value of *E_app_* was calculated for each pixel in a micro-spectroscopic image according to Eq. (3), for which a threshold criterion based on the calculated signal-to-noise ratio for both the donor and acceptor intensities in a given pixel (see Supplementary Methods section SM1) was then applied to the pixel-level maps of *E_app_*. This means that the background-subtracted intensity of both the donor and acceptor signal had to be greater than or equal to the desired threshold value, *TH*, times the standard deviation of the noise in a given pixel, or the corresponding *E_app_* value calculated for said pixel was rejected from further analysis. The *TH* was typically set to 1 for all experiments, except for a single experiment in which the wavelength channel bandwidth was set to 2 nm (see Section 2.2).

In certain pixels, the spectral unmixing resulted in an inadequate fit to the measured emission spectrum. In order to exclude these pixels from further analysis, we have introduced a second step of filtering, as follows. For every pixel within a micro-spectroscopic scan, the fit of the theoretical spectrum to the experimental one was quantified with the value *C*(*x,y*), which was computed according to Eq. (S2). For each segmented polygon (see section 3.1 and see Supplementary Methods section SM2), an average “goodness-of-fit” value was calculated by averaging the *C*(*x,y*) values found for each pixel within the segment. Individual segments with an average goodness-of-fit value which was greater than a fixed goodness-of-fit criterion were not considered for further analysis, as described in Supplementary Methods section SM3. If a segment did pass the goodness-of-fit criterion, the *E_app_* values from each of the pixels falling within the segment were then organized into a histogram plot, or a FRET spectrogram^28^, of bin width equal to 0.005. A FRET spectrogram was generated for each membrane segment in the set of images.

We further distilled the information contained within individual FRET spectrograms by extracting the positions of only the most prominent peaks within each spectrogram and generating a histogram of peak positions, i.e., a meta-histogram from the dominant peaks of multiple image segments^7,27,29^; the positions of the most prominent peaks were selected based on a routine described in detail in Supplementary Methods section SM4. The total number of peak positions used to generate each meta-histogram, *N_peaks_*, ranged from 100 to 300. The peak positions selected from individual FRET spectrograms were sorted based on the average receptor concentration (see section 2.4) of the corresponding segment such that meta-histograms were generated from peak values which originated from segments with the lowest overall concentration values.

As stated in the introduction, the meta-histogram is a collection of the *E_app_* values from the most probable FRET-productive oligomeric complexes and is the basis for determining the quaternary structure in FRET spectrometry. Each meta-histogram was modeled using an oligomeric structure with a particular size and shape^64^, as described in Supplementary Methods section SM5, in order to extract detailed information about the underlying receptor quaternary structure.

### 2.4 Estimation of receptor concentration for membrane cross-sections

The concentration of receptors within each membrane segment was obtained by calibrating the average donor and acceptor intensity values within the segment to fluorescence intensity values obtained from uniform solutions of known concentrations. Micro-spectroscopic measurements were performed on stock solutions of GFP_2_ and YFP fluorescent proteins at a number of concentrations, a process which has been previously described^27^. The same excitation wavelengths, power, and exposure times used to measure the yeast cells were applied to the fluorescent protein solution measurements. Calibration curves were generated for both the GFP_2_ and YFP solutions by plotting the average fluorescence intensity of the various solutions vs. the corresponding concentration. The slope, 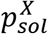, of the calibration plot represents the amount of fluorescence signal detected per molar concentration of fluorescent protein, and is proportional to the molecular brightness, 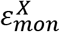, of a monomeric form of the respective fluorophore (see Eq. S11). The superscript *X* in the slope and molecular brightness symbols denotes either donor, *D*, or acceptor, *A*.

The method for converting fluorescence intensity values, obtained from fluorophore-tagged receptors residing in the plasma membrane, to receptor concentration is briefly presented below; for a more detailed description, see Supplementary Methods Section SM7.

The relation between the concentration of the *D* or *A* fluorophores in the plasma membrane, 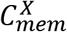, and the average measured intensity from the pixels within a given segment, 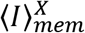, can be written as follows:

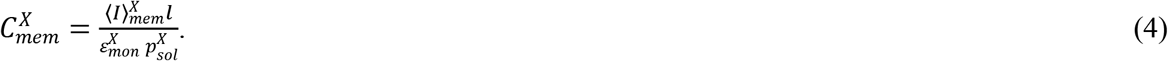

where *l* = 0.16μ*m* represents the length of a single camera pixel when projected onto the sample plane. To find 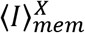 from the experimentally measured fluorescence values introduced in Eq.(2), we use the following relation:

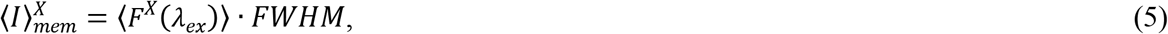

where 〈*F^X^*(*λ_ex_*)〉 is obtained by averaging the pixel level values of fluorescence of the donor in the absence of the acceptor (*F^D^*(*λ_ex_*)) or the fluorescence of the acceptor in the absence of donor (*F^A^*(*λ_ex_*)) within a single polygon segment. Because fluorescence emission from the membrane region is spread over multiple pixels of the detector, 〈*F^X^*(*λ_ex_*)〉 must be multiplied by *FWHM*, which represents the width in pixels of the emission *PSF* along the direction in which the signal is spread. According to Eq. (4), the average concentration (in molecules/μm^2^) of receptors tagged with a particular fluorophore (i.e., either *D* or *A*) within a given segment can be found by averaging the fluorescence intensity values of the corresponding fluorophore over all the pixels in the segment. Therefore, to find the total receptor concentration in a given segment, we must add the average concentrations of receptors tagged with both donor and acceptor:

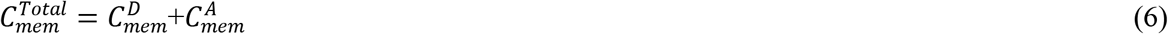

Using a simple relation between the measured fluorescence intensity and molecular brightness^65^, 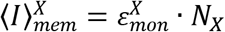, as well as the relation between molecular brightness and 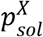 (Eq. S11), we can calculate the average number of donors or acceptors per pixel:

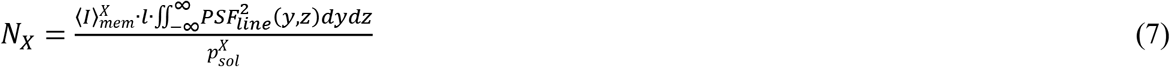

where *PSF_line_*(*y, z*) represents the spatial profile of the line shaped excitation beam, which is uniform along a single dimension (which we have denoted as the *x*-dimension in the theoretical formulation given in Supplementary section SM7).

In Eq. (7), we estimate the number of molecules contained within a volume corresponding to a single image pixel. This allows for the assembly of meta-histograms for specified concentration ranges, since the peaks chosen from *E_app_* histograms of segmented ROIs are calculated using numbers of image pixels.

## 3. RESULTS

### 3.1. Preliminary assessment of FRET efficiency meta-histograms

To probe the quaternary structure of the Ste2 receptor in living yeast cells (*S. cerevisiae*), we have implemented two-photon optical micro-spectroscopy^6,61^ to acquire pixel-level fluorescence spectra of yeast cells expressing Ste2-GFP_2_ and Ste2-YFP either singly or in combination with one another (see Materials and Methods). First, we obtained elementary emission spectra by imaging cells expressing only Ste2-GFP_2_, which was used as a donor of energy (D), or only Ste2-YFP, used as an acceptor (A), and then normalizing the intensity spectra to their maximum intensity values. Using the elementary spectra and a fitting algorithm described previously^62^, we unmixed the composite fluorescence spectra obtained for individual image pixels for cells co-expressing Ste2-GFP_2_ and Ste2-YFP. The unmixing procedure provided spatial intensity maps separately for the donor, *k^DA^*, and the acceptor, *k^AD^*, emission in the presence of each other (see Figures 1a and 1b). From such pairs of fluorescence maps, the apparent FRET efficiency, *E_app_*, was calculated^6,66^ for each image pixel (see Figure 1c).

**Figure 1.**
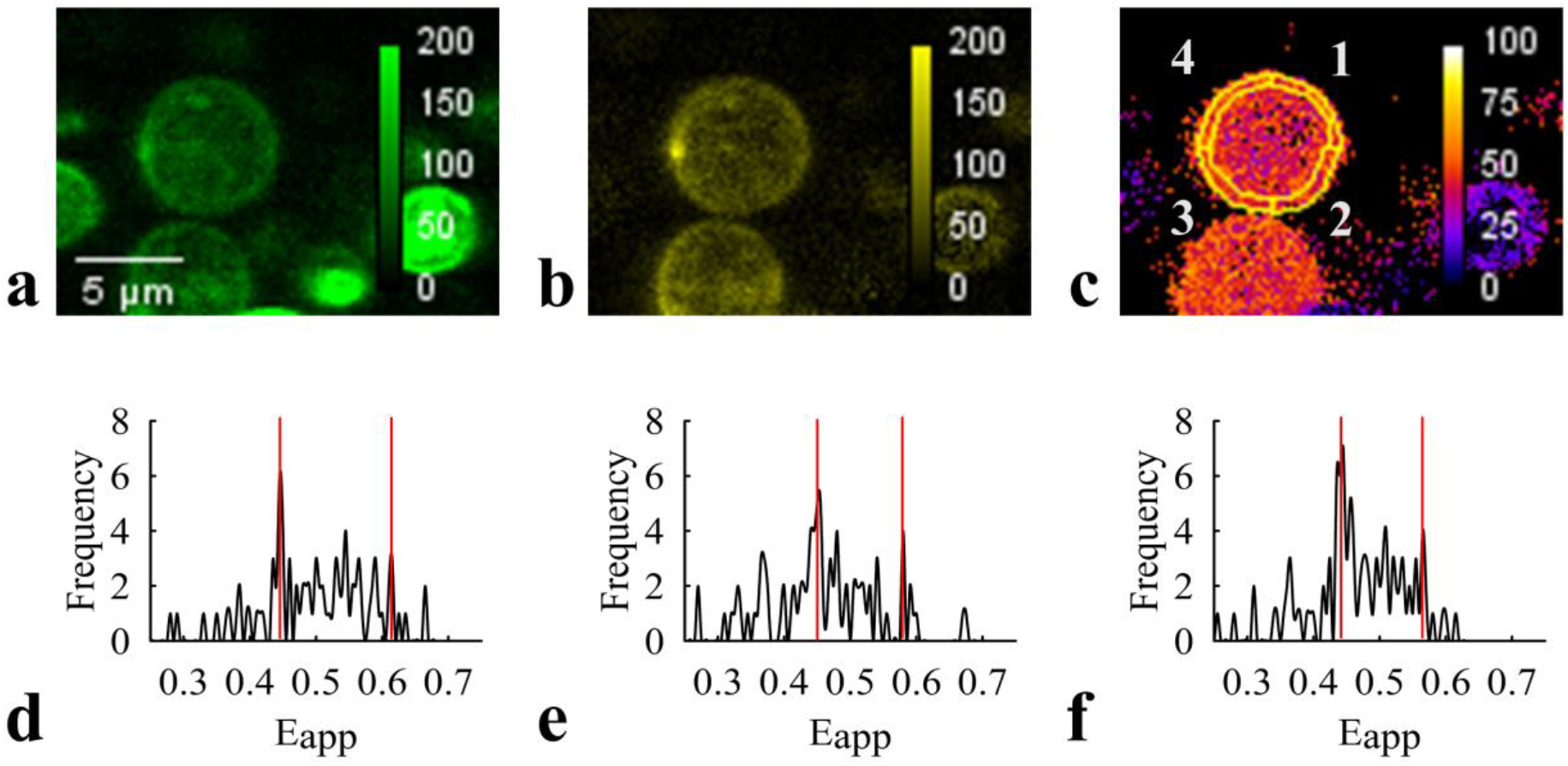
Typical results obtained from imaging yeast (*S. cerevisiae*) cells co-expressing Ste2-GFP2 and Ste2-YFP. Spectral unmixing provided separate maps of the fluorescence signals of (a) donors in the presence of acceptors, *k^DA^* and (b) acceptors in the presence of donors, *k^AD^*. (c) Apparent FRET efficiency, *E_app_*, maps were determined from the pixel-level values of *k^DA^* and *k^AD^*, as described in the Materials and Methods section. Contours defining regions of interest (ROI) were hand-drawn around the exterior of cells, and pixels were removed from the interior of the selections until a ring (with a width of ten pixels) encompassed only the plasma membrane; each such ROI was split into four segments as described in the supplementary methods section SM2 and Figure S1. Segmented ROIs were transferred to the *E_app_* maps (shown as yellow curves in panel c), and histograms showing the number of pixels within a certain *E_app_* bin range (Frequency) for different *E_app_* values were generated from each segment using a bin size of 0.005 for *E_app_*. Representative *E_app_* histograms are shown in panels d, e, and f from segments 1, 3, and 4, respectively. The positions of the two most dominant peaks of these histograms (indicated with vertical red lines) were among the ones used, as described in the Materials and Methods, to generate the meta-histograms shown in Figure 2.

Hand-drawn polygonal regions of interest (ROI) were made for each cell within either the *k^DA^* or *k^AD^* maps (whichever had a more clearly defined outline), and pixels belonging primarily to the cytoplasm were then removed from the maps using a computer algorithm that only retained a band ten-pixel wide within the ROI, as measured from the exterior of the ROI (see Supplementary Section SM2 for a full description of the method). The resulting polygonal rings were subsequently divided into four segments using the computer program. The segmented ROIs were then transferred to the *E_app_* maps (see Figure 1c for typical results), from which FRET efficiency histograms were generated (Figure 1d-f) by binning together the pixels with similar FRET efficiencies at a bin interval of 0.005. The *E_app_* histograms from the randomly selected cell displayed in Figure 1 revealed varying levels of detail (i.e., locations along the *E_app_* axis, amplitudes, and widths of the peaks in the histograms) that are often indicative of the presence of receptor oligomers, as previously described^27,29^.

In order to determine the most probable quaternary structure of Ste2 oligomers in the absence and presence of the α-factor pheromone, we exploited the large number of histograms obtained to generate ensembles of the most abundant configurations of the oligomeric structures. To this end, we extracted specific *E_app_* values corresponding to two clearly visible peaks from each histogram (indicated by vertical red lines in Figure 1, panels d, e, f) using an algorithm described in Supplementary Methods section SM3. We used these peak values to generate a histogram of peak positions, which is called a *meta-histogram*^6–7,29,62,67^. An *E_app_* bin width of 0.02 was used to assemble the meta-histograms.

As done previously^27^, we selected only ROI segments with relatively low receptor expression levels in the assembly of *E_app_* meta-histograms, in order to avoid the use of featureless histograms generated by mixing of multiple configurations of oligomers in pixels where the concentration of receptors is very high. We found that a good compromise between meta-histogram resolution and number of experimental data points occurred when choosing total receptor concentrations ranging from ~2 (corresponding to the lowest receptor expression level) to 43 receptors per pixel^2^ (with an average of 19 receptors per pixel), which corresponds to a concentration range of ~9 to 211 molecules/μm^2^ (an average of 96 molecules/μm^2^).

To determine average concentrations of receptors within each membrane segment, which was needed to filter the segment-level *E_app_* histograms used to assemble the meta-histograms, we first scanned the sample with 930-nm laser light (which provides good acceptor excitation) and then with 800-nm light (which only excited the donor). The 800-nm excitation was used to extract the apparent FRET efficiency, while both excitation wavelengths provide measured intensities used to estimate molecular concentrations of both the donors and acceptors^27^. Because photo-switching and/or photobleaching of the donor may occur more appreciably upon excitation at 800 nm, the order of excitation scans (i.e., first 930 nm and then 800 nm) was chosen to minimize donor photo-switching and/or photo-bleaching prior to excitation at 930 nm^68–70^. Conversely, minimal changes in the fluorescence intensities obtained upon excitation at 800 nm are observed when the samples are first excited at 930 nm.

Examples of *E_app_* meta-histograms obtained for two different ranges of receptor concentrations (i.e., 40.7 < molecules/μm^2^ < 132.0 and 40.7 < molecules/μm^2^ < 159.6) are shown in Figure 2a-d. For each concentration range, the meta-histograms were assembled by choosing either a single peak (panels a and c) or two peaks (panels b and d) per *E_app_* histogram, for comparison.

**Figure 2.**
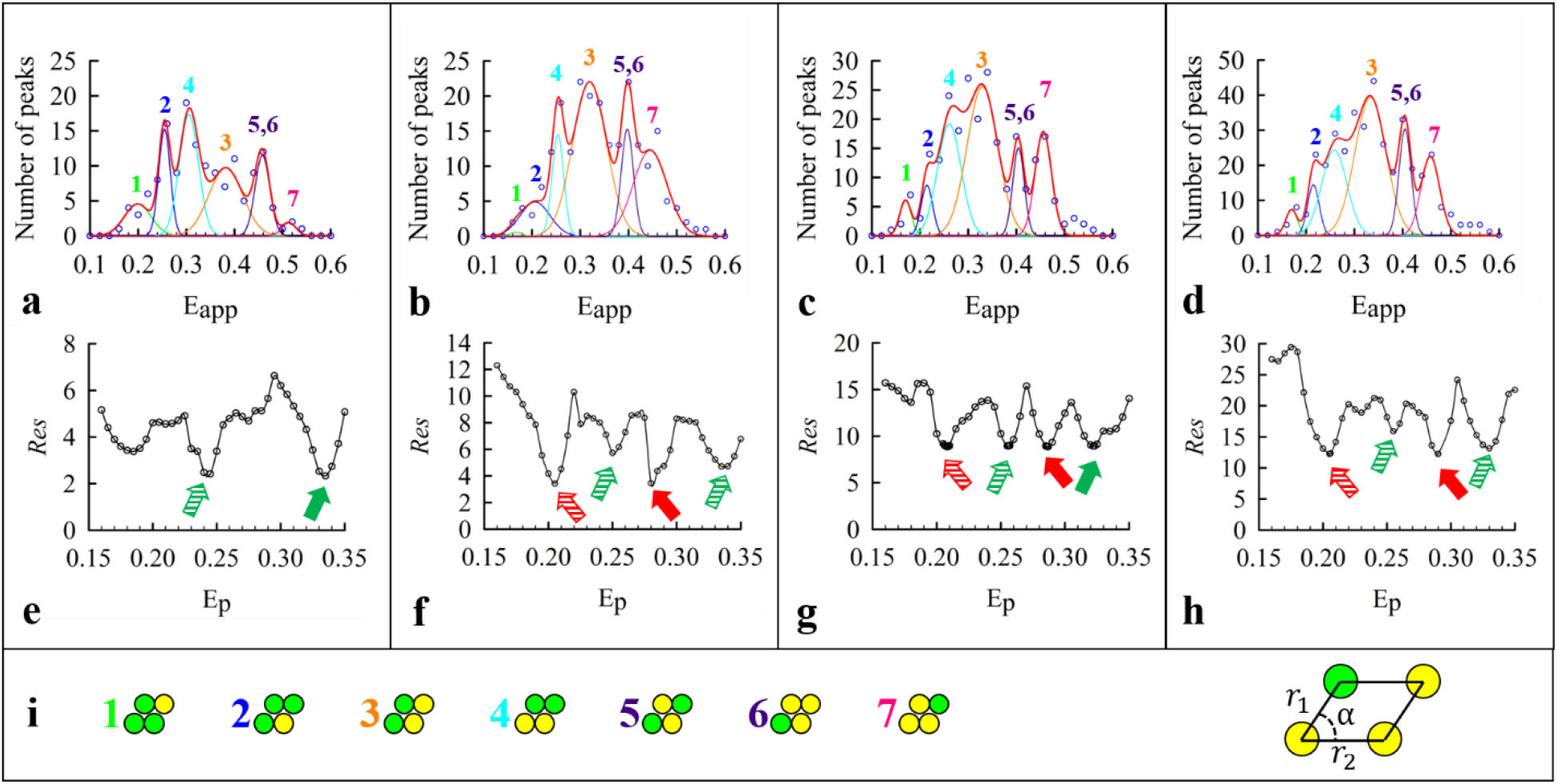
Typical meta-histograms obtained from yeast (*S. cerevisiae*) cells co-expressing Ste2-GFP2 and Ste2-YFP in the absence of α-factor at low receptor concentrations, and their analysis using an appropriate quaternary structure model. Experimental meta-histograms (empty blue circles) obtained by collecting either single peaks (panels a and c) or two peaks (panels b and d) from each *E_app_* histogram of the type shown in Figure 1 were fitted (solid red lines) using a reduced residual minimization algorithm (see section SM5 and panels e-h) to a general parallelogram-shaped tetramer model, which is comprised of seven Gaussian peaks corresponding to particular FRET-productive configurations of donors and acceptors within a tetramer (see panel i). The meta-histograms were assembled for two different receptor concentration ranges: 41 to 132 receptors per μm^2^ (panels a and b) and 41 to 160 receptors per μm^2^ (panels c and d). Positions of the seven meta-histogram peaks predicted by the model depend on only three parameters, *E_p_* (pairwise FRET efficiency), *r*_1_/*r*_2_, and angle *α* (shown in i), which are used as adjustable parameters in the data fitting process. At the beginning of the fitting process, *E_p_* was first set to 0.16, and the lowest possible reduced fitting residual (see Supplementary Methods section SM5) was obtained by adjusting the fitting parameters. The process was repeated several times after increasing *E_p_* in a stepwise manner, and the reduced fitting residual vs. *E_p_* were plotted (panels e-h). One such curve computed for each histogram is shown under its respective meta-histogram in the figure. Note that there is a pair of complementary fits for each curve, indicated by two arrows with the same color, and which are obtained by simply switching between *r*_1_ and *r*_2_ values in the model. Since the two situations are structurally indistinguishable, only one of the fits (indicated by solid color arrows) is retained in our subsequent analysis. In addition, the *Res* vs. *E_p_* plot may present two sets of such local minima, each set denoted by arrows of different colors. Most of the time, only one set of complementary fits provides the global minimum, but sometimes two such sets take similarly low *Res* values. Best-fit parameters and *Res* corresponding to the minimum indicated by the solid color arrow(s) were: *E_p_*= 0.335, *r*_1_/*r*_2_ = 0.93, *α* = 66.33, and *Res* = 2.33 for panels (a) and (e); *E_p_*= 0.280, *r*_1_/*r*_2_ = 0.93, *α* = 68.27, and *Res* = 3.44 for (b) and (f); *E_p_*= 0.287 and 0.323, *r*_1_/*r*_2_ = 0.93 and 0.95, *α* = 67.23 and 59.99, and *Res* = 8.85 and 8.90, for (c) and (g); *E_p_*= 0.290, *r*_1_/*r*_2_ = 0.93, *α* = 66.95, and *Res* = 12.24 for (d) and (h).

Peaks observed in meta-histograms have previously been ascribed to the FRET-productive configurations of the most probable receptor quaternary structure^27–29^ and have been used to extract geometrical parameters corresponding to quaternary structure models^6–7,29,62,67^. The quaternary structure models predict a number of Gaussian peaks, whose relative positions (i.e., “means”) are determined by only three parameters, the pairwise FRET efficiency, *E_p_*, the ratio of the side lengths of the oligomer, *r*_1_/*r*_2_, and the acute angle between the sides, *α* (see Figure 2). Inspired by the previous analyses published by our lab involving Ste2^6,27^, we first analyzed our current meta-histograms, constructed from segments with low average receptor concentrations (i.e., 2 to 25 receptors/pixel or 11.2 to 140 receptors per μm^2^), using a rhombus-shaped (i.e. *r*_1_/*r*_2_ = 1) tetramer model with an acute angle between sides of the rhombus of *α* = 60°. However, it has been determined that a parallelogram-shaped tetramer model better estimates the shape of the quaternary structure for a number of other GPCRs^7,29^. Therefore, because of the large amount of high-quality data collected in this study, we wanted to refine the modeling of the Ste2 quaternary structure to see whether a more general parallelogram shape better approximates the structure, as compared to the previously used rhombus shape. In addition to the FRET determining parameters, the theoretical curves predicted for each model also include the amplitudes of the individual Gaussians, which depend directly on the frequencies of occurrence of each FRET-productive configuration^6,28^, and the width of the Gaussians, which depends on the relative angles between the transition dipoles of each fluorescent tag and the degree of cylindrical averaging taking place during measurements^6,71^ but which is not specifically included in our theoretical model.

The Gaussian amplitudes, standard deviations, and the parameters determining the Gaussian peak positions were adjusted systematically for each model used, in order to minimize the fitting residual, i.e., the sum of the squared differences between experimental and theoretically predicted data points. A comparison between the results obtained with the rhombus and general parallelogram models was quantified using a reduced fitting residual, given by Eq. (S6) of Supplementary Methods Section SM5. The reduced fitting residual, *Res*, was computed by dividing the fitting residual by the number of degrees of freedom of the data, i.e., number of data points minus number of fitting parameters. The reduced fitting residual was needed to properly compare the two models, due to the fact that the general parallelogram model has more fitting parameters than the rhombus tetramer model (*r*_1_/*r*_2_ = 1 and *α* = 60° were held fixed for the rhombus model as stated above). The general parallelogram-shaped tetramer model fitted the data significantly better than did the rhombus tetramer model, as seen by the plots shown in Supplementary Figure S4 and the vastly lower reduced residuals (i.e., 3.048 for parallelogram vs. 7.836 for rhombus in absence of ligand, and 0.938 for parallelogram vs. 7.504 for rhombus in the presence of ligand).

While the deviations of the quaternary structure model of Ste2 from that of a rhombus revealed by this preliminary analysis were relatively small (i.e., less than 10% difference between the lengths of the two sides of the parallelogram, with the angle between them just a few degrees larger than 60°), being able to capture them with such exquisite precision allows us to ask more detailed questions regarding the possible effect of ligand binding on the geometry of the oligomer. This ability to extract small differences in the quaternary structure of Ste2 performed in the present study is due to several modifications to the method as discussed in the introduction. Furthermore, our current ability to automatically separate membrane regions within the images into smaller segments allowed the capture of local fluctuations in FRET efficiencies and permitted generation of larger pools of data (~3,000 cells/experiment) for dramatically improved statistics. Based on the results of this preliminary analysis, we decided to apply the parallelogram model to all meta-histograms in the absence and presence of ligand going forward.

For typical results pertaining to samples treated with the α-factor ligand and analyzed in a similar manner as shown in Figure 2, see Supplementary Figure S5.

### 3.2 Computer simulations reveal the effect of noise on meta-histograms

One of the first questions that may arise while glancing at the meta-histograms of the types displayed in Figure 2 concerns the extent to which noise affects the visibility of the various peaks, and the ability to extract oligomer geometrical information from their number and location. To address this potential concern, we have conducted computer-based numerical simulations at the level of single oligomeric complexes per pixel, with an aim to replicate conditions leading to experimental meta-histograms (see Supplementary Methods sections SM8 and SM9 for a detailed description of the simulations). For simplicity, in these simulations we opted to use the rhombus-shaped tetramer model for the oligomeric complex to be simulated. Using *E_p_* = 0.20, *α* = 60°, and *r*_1_/*r*_2_ = 1, we calculated the theoretical *E_app_* peak positions, along with the expected donor and acceptor intensity levels, for all configurations of the rhombus. Using the expected donor and acceptor intensities for a given configuration, we created normal probability distributions, defined by Eq. (S23), which were centered at the expected intensity values of each. The width of the normal probability distributions was determined from a fixed signal-to-noise ratio (*SNR*), according to Eq. (S24). For each pixel in a simulation, a random intensity value for both the donor and acceptor was chosen from the corresponding probability distribution which was constructed for the particular oligomer configuration found in said pixel, and a value of *E_app_* was then calculated by plugging the two randomly chosen intensity values into Eq. (3). *E_app_* histograms were constructed from a collection of 1000 pixels, and a meta-histogram generated from 500 *E_app_* histograms, using the same protocol described in section 2.3. Since the signal level in all simulations was fixed to intensity values which correspond to a single tetramer for each simulated pixel, we could generate meta-histograms for a wide range of signal-to-noise ratios (*SNR*) simply by changing the width of the normal probability distributions constructed using Eq. (S23).

Although a naïve prediction would be that noise may generate artificial peaks in the meta-histograms, our simulations showed that addition of large amounts of noise generated no additional peaks. For very low values of *SNR* (≤ 1), the meta-histograms did contain a smooth broad background distribution across all *E_app_* values which was centered around *E_app_* = 0.5 (see Supplementary Figures S9 and S10). However, within these histograms generated for *SNR* ≤ 1, individual peaks, while relatively small, are still clearly visible, and their locations corresponded to the FRET values of the various tetramer configurations in the simulation. The range of *SNR* levels corresponding to live-cell experiments were determined based on a comparison of live-cell intensity measurements to those from a control experiment involving characterization of EMCCD noise for various light intensity levels (see Supplementary Methods section SM9). The simulated meta-histograms with *SNR* levels corresponding to our live-cell experiments, i.e., 20 ≤ *SNR* ≤ 30 on average, are indicated in Supplementary Figures S9 and S10 with red boxes. Thus, from our simulated meta-histograms, we can safely assume that noise does not generate artificial peaks in the experimental meta-histograms and that in fact the challenge is to reduce the noise such that the peaks are not smeared or even obliterated.

### 3.3 Detailed meta-histogram analysis reveals multiple oligomeric conformation sub-states

We employed an iterative process for fitting the meta-histograms with the parallelogram shaped tetramer model. In this fitting procedure, one parameter, *E_p_*, was fixed at a particular value, and all other parameters were adjusted until the reduced fitting residual, *Res*, was minimized. Then the value of *E_p_* was increased by a small increment, and the fitting procedure involving all other parameters run again. This iterative fitting procedure was repeated for a large range of *E_p_* values (0.16 ≤ *E_p_* ≤ 0.5), and plots of the minimum *Res* value obtained for each *E_p_* value were plotted against the corresponding *E_p_* value; examples of *Res* vs. *E_p_* plots are shown in Figures 2e-h.

As seen in Figure 2, fitting the *E_app_* meta-histograms with a mathematical expression generated from the parallelogram-shaped tetramer for a range of *E_p_* values leads to one global minimum and possibly a second, local, minimum in plots of *Res* vs. *E_p_* (after excluding complementary minima corresponding to swapping between *r*_1_ and *r*_2_ values). If data from a few extra membrane segments were added to the original meta-histogram, the best-fit theoretical meta-histogram peaks sometimes shifted positions slightly, which corresponded to the local minimum becoming a global minimum and the global minimum taking its place as a local minimum. As we added peaks from additional segments to the meta-histogram, without significantly increasing the upper limit on the concentration range from which the segments were selected, the minimum *Res* value oscillated between the two different *E_p_* positions. We took this as evidence that, in fact, there is more than one quaternary structure sub-state present in the system, each characterized by its own set of *E_p_*, *r*_1_, *r*_2_, and *α* values and corresponding to a certain local minimum in the *Res* vs. *E_p_* plots. Accordingly, a set of segment-level histogram peaks may lead to a meta-histogram in which one quaternary structure sub-state dominates. However, for another set, another sub-state may dominate, just by pure chance.

To investigate this hypothesis systematically, we first took all 2,332 (ligand, absent) and 2,370 (ligand, present) histogram peaks obtained from segments with receptor concentrations less than 211 molecules/μm^2^, separated them into groups based on the experimental day they were acquired, and then listed each group in ascending order of their total receptor concentration. We then created *E_app_* meta-histograms for each experimental day using the first 100 histogram peaks in the sorted list (i.e. peaks which originated from the segments with the lowest concentration values). These initial meta-histograms were then analyzed using the iterative fitting procedure described above and illustrated in Figure 2. Next, we created additional meta-histograms by adding peaks, typically in increments of 50, to the existing ones. This process of building up meta-histograms incrementally was repeated until the meta-histograms for a particular experimental day contained 300 peaks; the highest concentration of a segment used in any of the constructed meta-histograms was 211 receptors/μm^2^. In special cases where we noticed a switch from a single minimum to two minima along the *Res* vs. *E_p_* curves, e.g., as shown in Figure 2e-h, we occasionally introduced a smaller increment of 25 peaks in an attempt to capture both minima.

Meta-histograms were analyzed using the iterative fitting procedure for each incremental step in number of peaks; a total of 60 meta-histograms were constructed and analyzed for data obtained in the absence of ligand, and 64 in the presence of ligand. From the different sets of best-fit *E_p_* and *r*_1_/*r*_2_ values obtained for the quaternary structure in the absence and presence of α-factor ligand, the two parallelogram side lengths were computed using the expression *r* = *R*_0_(1/*E_p_* − 1)^1/6^. In this regard, the previously published value of the Forster radius for the GFP_2_/YFP pair, *R*_0_= 57 Å^55^, serves as a “molecular ruler”^72^. The 60 sets of *r*_1_ and *r*_2_ distances thus obtained in the absence of ligand and 64 in the presence of ligand were used to generate separate histograms (referred to as *r*_1_ or *r*_2_ distance histograms) for the frequency of occurrence of *r*_1_ and *r*_2_ distances (see open black circles in Figure 3). At this exquisite (sub-Ångstrom) resolution, one can clearly distinguish two or three peaks in the distributions of distances, indicating the presence of multiple quaternary conformation sub-states (see Discussion section below for elaboration on this point).

**Figure 3.**
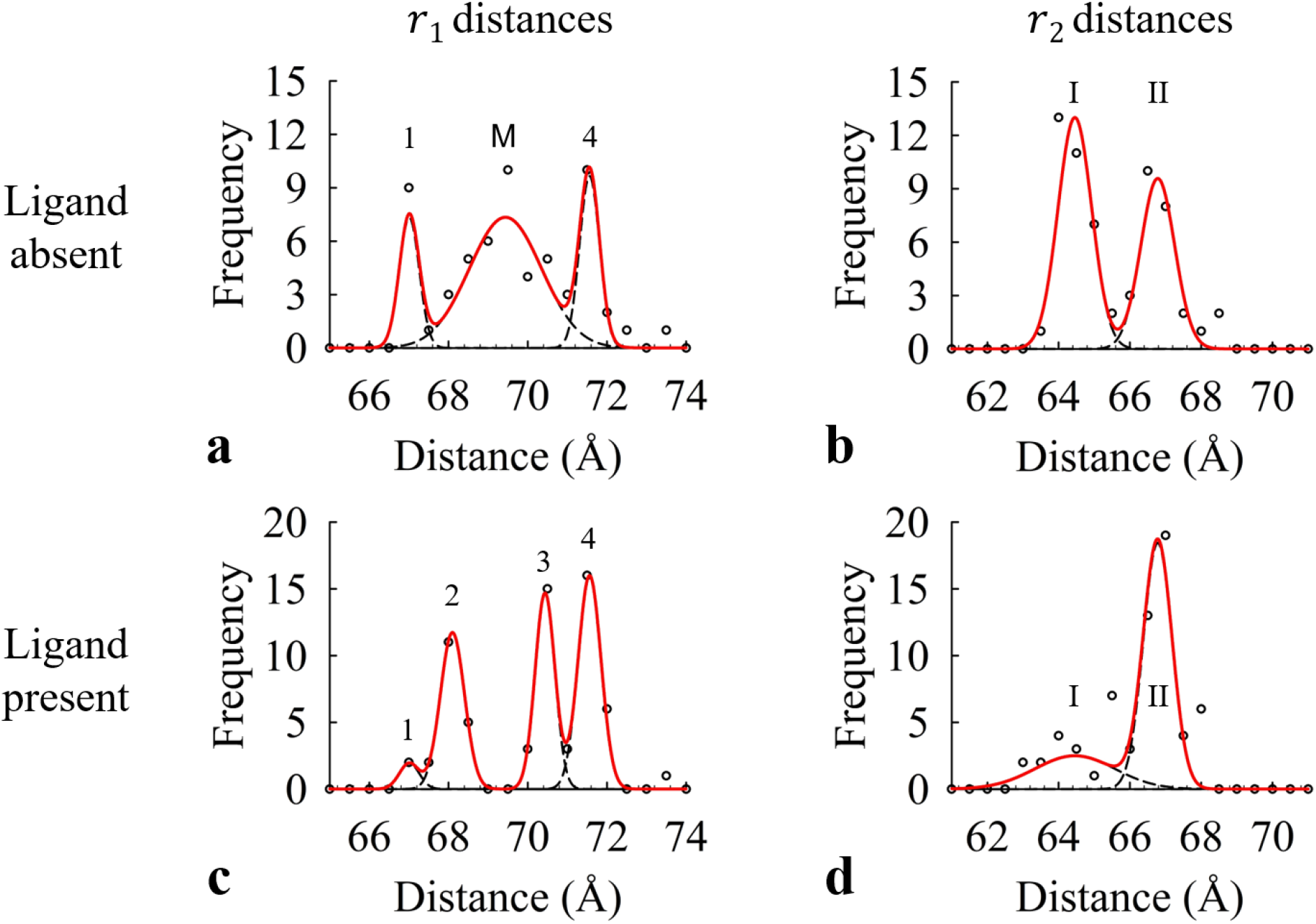
Histograms of frequencies of occurrence of the side lengths of the general parallelogram-shaped tetramer obtained from fitting the theoretical model (Figure 2) to the experimental meta-histograms. (a) The *r*_1_ distance values (black circles) obtained in the absence of α-factor were binned into 0.5 Å bins and fit with a model consisting of a sum of three Gaussian functions (dashed black lines) corresponding to three different conformational states. The Gaussian parameters were adjusted to minimize the reduced *χ*^2^ between the fitted model curve (solid red line) and the experimental data points. (b) Similarly, the *r*_2_ distance values in the absence of α-factor were used to generate a histogram (with 0.5 Å bin size) which was fit with a sum of Gaussian curves corresponding to two conformational states found for this distance. The same protocol described for panel a was used to minimize the reduced *χ*^2^ between the fitted curve and the experimental points. (c) Histograms for *r*_1_ distance values obtained in the presence of ligand and fit with a sum of four Gaussians. (d) Histograms for *r*_2_ distance values obtained in the presence of ligand and fit with a sum of two Gaussians. Plots in panels a and b each contain 60 data points, while those in panels c and d contain 64 data points (corresponding to as many meta-histograms of FRET efficiencies). The best-fit parameter values of each of the Gaussians used to fit the distance histograms are listed in Tables 1 and 2.

The positions of the different peaks in the *r*_1_ and *r*_2_ histograms were determined by fitting a sum of Gaussian functions (represented by solid red lines in all panels of Figure 3) to the experimental data points (shown as empty circles). Since some of the peaks (labeled by “1” and “4,” in the *r*_1_ histogram and by “I” and “II” in the *r*_2_ histogram) were visible in both the presence and absence of ligand (albeit with different amplitudes), the fitting required that the mean positions of those Gaussian functions were held fixed relative to one another during the process (i.e., simultaneously fit). However, when comparing the distributions of the *r*_1_ data for ligand treated vs. ligand absent, the peaks located between peak 1 and 4 do not align in a similar fashion. Therefore, a single Gaussian function was used to fit the middle peak of the *r*_1_ histogram (labeled by “M” for mixture) in the absence of ligand, whereas two separate Gaussian functions were used to fit the peaks labeled “2” and “3” in the *r*_1_ histogram obtained in the presence of ligand.

From the fitting of the *r*_1_ distance histogram in the presence of α-factor, we can identify the two middle peaks in Figure 3c (i.e., peak 2 and 3) as intermediate conformational states between the two fixed peaks (i.e., peak 1 and 4) since they do not exist in the absence of α-factor. As is clearly seen in Figures 3b and d, the relative amplitude of peaks I and II of the *r*_2_ distance histograms change when ligand is added, with the peak II amplitude increasing and, thereby, the peak I amplitude decreasing. To quantify such changes in the relative abundance of all the distance histogram peaks when ligand was added, we calculated the relative percentage fraction for each peak by evaluating the area under each Gaussian function and then dividing each by the total sum of the area under all Gaussians used in the model (see Figure 4).

**Figure 4.**
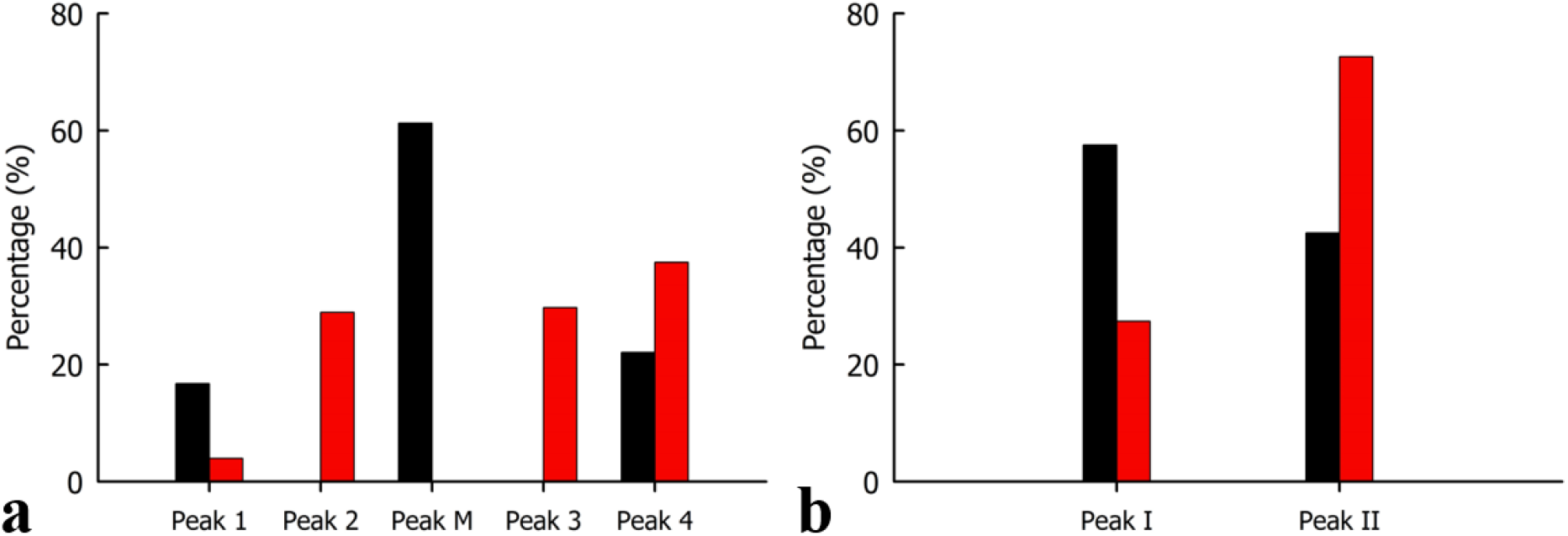
Relative abundance of the conformational states for *r*_1_ and *r*_2_ distances of the parallelogram-shaped tetramer formed by Ste2 in the absence and presence of α-factor. (a) To compute Percentage (%), the area under each Gaussian used to fit the *r*_1_ distance histograms shown in Figure 3a and 3c is divided by the total area under the curve of their respective models (i.e., the sum of the Gaussian curves comprising the model). In the absence of α-factor (black bars), three Gaussian curves (or “peaks”) are required to best describe the data (labeled here as peak 1, M, and 4), while in the presence of α-factor, four distinct peaks occur (described by peaks 1, 2, 3, 4). The Percentage (%), or relative abundance, of each peak within a specified model shows which states are more favored in each scenario. (b) Similarly, the relative abundance of each Gaussian curve used to model the *r*_2_ distance histograms of Figure 3b and 3d were computed using the protocol described in panel a, in both the absence and presence of α-factor (black and red bars, respectively). Here, the model is comprised of two Gaussian curves (I and II), for which the amplitudes of the bars show a major shift from absence to presence of α-factor, indicating preference of conformational states under each scenario.

The bar chart of the relative abundance of *r*_1_ distance values presented in Figure 4a shows that in the absence of ligand, peak M (*r*_1_=69.4 Å) dominates the distribution, but when ligand is present peaks 2 (*r*_1_=68.1 Å) and 3 (*r*_1_=70.4 Å) are nearly equal in amplitude (29% and 30%, respectively) with peak M disappearing entirely. Peak 1 (*r*_1_=67.0 Å) decreased while peak 4 (71.6 Å) increased, making the longer *r*_1_ distances more probable than the shorter ones in the presence of ligand (see amplitudes and standard deviation values in Table 1). Similarly, the relative abundance of *r*_2_ distance values (Figure 4b) changed more markedly after ligand was added, with the peak corresponding to a longer *r*_2_ distance increasing appreciably. Specifically, Peak I (*r*_2_=64.5 Å) became significantly less abundant than peak II (*r*_2_=66.8 Å) in the presence of ligand, indicating a significant propensity of the structure to assume conformations with longer distances between protomers (i.e., 66.4 Å≤ *r*_2_ ≤ 67.2 Å; see amplitudes and standard deviations in Table 2).

**Table 1.**
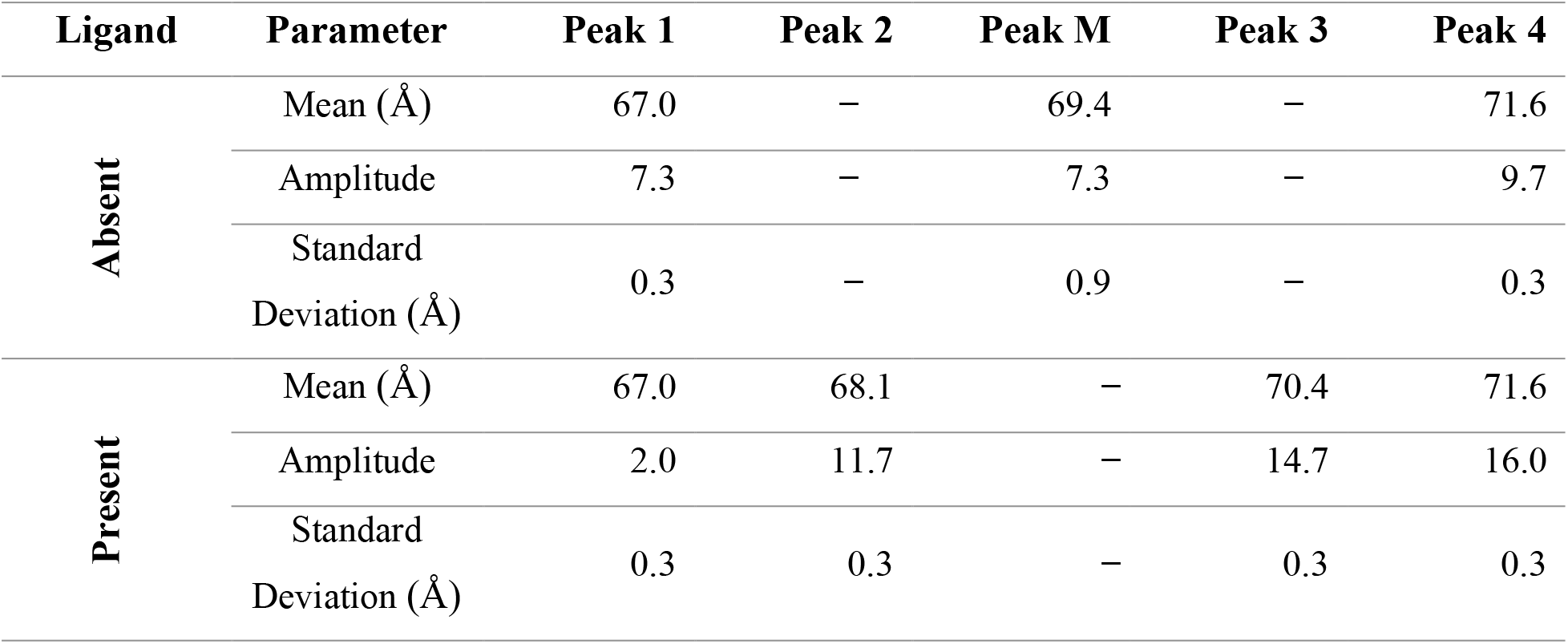
Best-fit parameters for the multiple Gaussian curves used to model the *r*_1_ distances found for the various conformational states of the parallelogram-shaped tetramer for Ste2 shown in Figures 3a and 3c.^1^

| Ligand | Parameter | Peak 1 | Peak 2 | Peak M | Peak 3 | Peak 4 |
| --- | --- | --- | --- | --- | --- | --- |
| Absent | Mean (Å) | 67.0 | – | 69.4 | – | 71.6 |
|  | Amplitude | 7.3 | – | 7.3 | – | 9.7 |
|  | Standard Deviation (Å) | 0.3 | – | 0.9 | – | 0.3 |
| Present | Mean (Å) | 67.0 | 68.1 | – | 70.4 | 71.6 |
|  | Amplitude | 2.0 | 11.7 | – | 14.7 | 16.0 |
|  | Standard Deviation (Å) | 0.3 | 0.3 | – | 0.3 | 0.3 |

**Table 2.** Best-fit parameters for two Gaussians used to model the distributions of *r*_2_ distances found for the conformational states of the parallelogram-shaped tetramer for Ste2 shown in Figures 3b and 3d.^2^

| Ligand | Parameter | Peak I | Peak II |
| --- | --- | --- | --- |
| Absent | Mean (Å) | 64.5 | 66.8 |
|  | Amplitude | 13.0 | 9.6 |
|  | Standard Deviation (Å) | 0.5 | 0.5 |
| Present | Mean (Å) | 64.5 | 66.8 |
|  | Amplitude | 2.5 | 18.4 |
|  | Standard Deviation (Å) | 1.1 | 0.4 |

## 4. DISCUSSION

Conformational changes occurring to the tertiary structure of GPCRs during the activation process has been heavily investigated in recent years. Early hypotheses regarding possible GPCR tertiary structures depicted the receptor as a simple switch which existed in either an inactive or active state. However, a growing body of evidence suggests that activation of GPCRs is not a binary operation, but rather that the tertiary structure conformation can exist in a series of intermediate states^50^, with a range of activity levels possible across the distribution of conformations as shown by fluorescence-based techniques^30,73^, FNMR^32^, AFM^74^, electron microscopy^75^, X-ray crystallographic studies^38^, and theoretical simulations using crystal structures^2,33,76–77^. From crystal structure studies of individual GPCRs, it has been observed that the intracellular region of TM1, TM3, TM5, TM6, and TM7 move outward from one another when ligand is bound^2^, with TM6 showing the most pronounced outward movement in the cytoplasmic end of the helix as compared to its membrane-integrated portion, and TM5 showing helical extension^38,77^.

The purpose of our study was to investigate the quaternary structure organization of the Ste2 receptor, with particular emphasis on what effect, if any, ligand binding has on this structure. As it will be discussed in more detail below, the results presented in Section 3 strongly indicate that not only the tertiary structure of individual GPCR protomers within a protein complex but also the quaternary structure may exist in multiple sub-states; in fact, there must be a causal relationship between the two, as hinged motions of transmembrane domains belonging to abutting protomers may result in changes in the distance between the center of mass of the protomers. Therefore, monitoring changes in quaternary structure in living cells allows one to extract information on changes in the tertiary structure.

In order to help visualize the likelihood of the Ste2 quaternary structure to be in a particular substate and furthermore illustrate how ligand binding affects the probability associated with the occupancy of each such substate^30,32,73^, we have drawn theoretical potential energy landscapes^30,32–33,50,73^ depicting the basal and active conformations of the oligomer, as seen in Figure 5. Potential energy landscapes, which have been extensively used as a convenient visual tool for discussing protein conformations, represent the energy of the quaternary conformations along the receptor activation pathway^30,32–33,50,73^. Conformational states with lower energy are more stable, and thus are more likely to be populated. These stable conformations are represented by potential wells, or minima, in the energy landscape. The number and depth of such wells illustrated in Figure 5 were informed by analysis of the *r*_1_ and *r*_2_ histograms of Figure 3 and relative abundance plots of Figure 4. Specifically, the relative abundance of a particular stable quaternary conformational state is reflected in the potential energy landscape by the depth of the associated well, while the likelihood of transitioning from one state to another is reflected in the height of the energy barrier between them.

**Figure 5.**
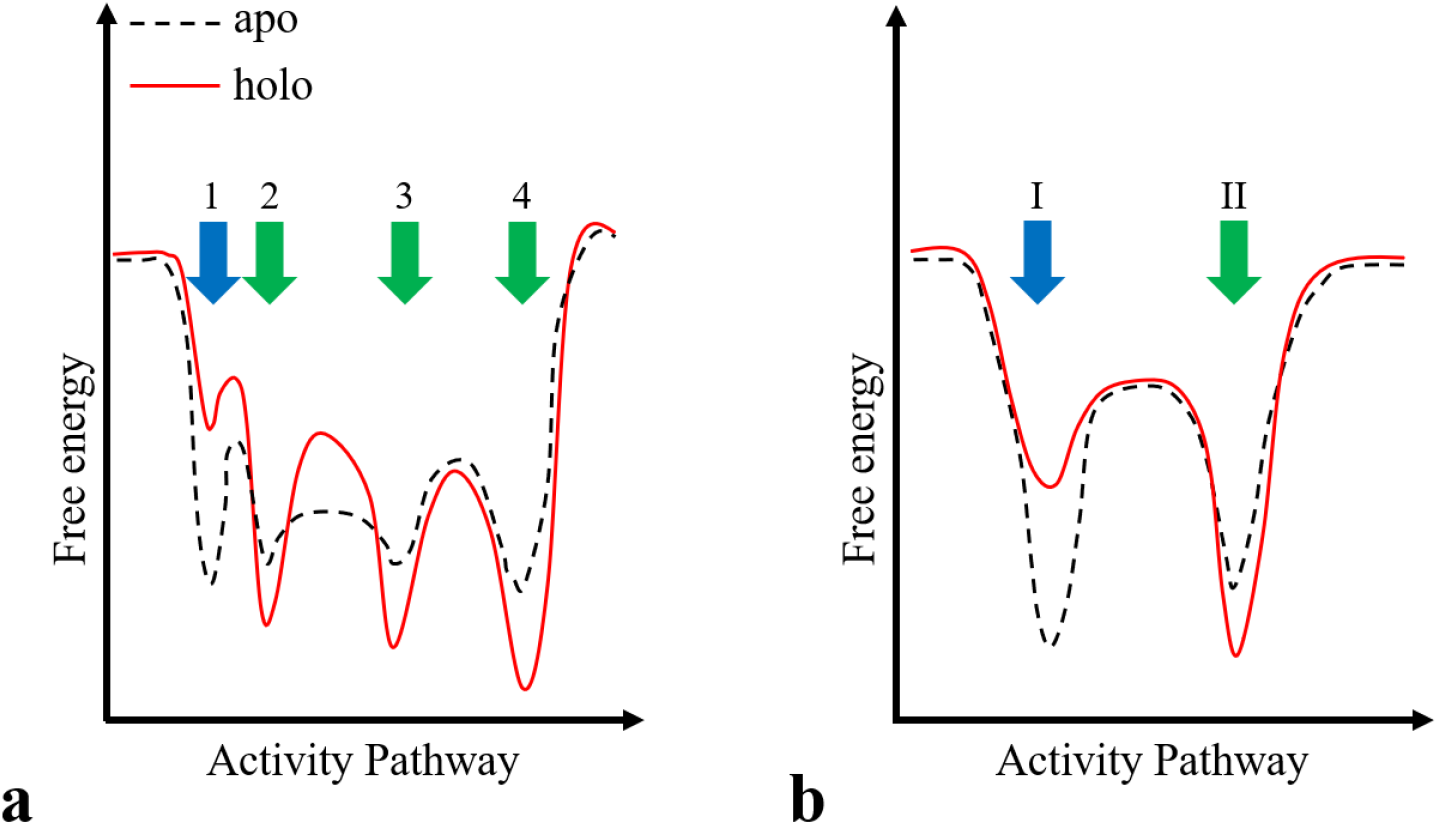
Schematic representation, using free energy landscape diagrams, of the multiple conformational states of the Ste2 quaternary structure in absence and presence of α-factor pheromone corresponding to the *r*_1_ (a) and *r*_2_ (b) distances between protomers within an oligomer. The schematics were drawn based on the distributions of the *r*_1_ and *r*_2_ side lengths of the parallelogram representing the Ste2 oligomer shown in Figure 3 and their relative abundance presented in Figure 4. The *r*_1_ distribution obtained in the absence of ligand shows how none of the four states is more probable than the next, and the barrier between active state 2 and active state 3 is lower than the barrier between other states, making these states more unstable than the inactive basal state 1 and active state 4. When ligand is present (solid red curve), state 1 diminishes while the probability of the receptor residing in the higher activity states (2, 3, 4) increases, as reflected by the lowering of energy minima upon binding of ligand. For the free energy diagram corresponding to *r*_2_ distances, both the inactive (I) and active (II) conformations are present in abundance in the absence of ligand, with a slight preference for the inactive state. When ligand is administered and bound to receptors within the oligomer, the active conformation dominates.

By examining the *r*_1_ and *r*_2_ distance histograms of Figure 3 and the relative abundances of the Ste2 conformations shown in Figure 4, two significant observations may be made, which enable us to generate a picture describing the effect of ligand on the probability associated with the occupancy of each quaternary conformation. Firstly, we see in Figure 4 that treatment of Ste2 with α-factor altered the relative abundance of each of the quaternary conformational states when compared to the conformations of Ste2 in the absence of ligand except for peaks 2, 3, and M in Figure 4a. Specifically, the relative abundance of the shortest *r*_1_ and *r*_2_ distances (labeled peaks 1 and I in Figure 4) both decreased appreciably when ligand was administered. Conversely, the abundances of the states characterized by longer *r*_1_ and *r*_2_ distances (labeled peaks 4 and II in Fig. 4) both increased upon addition of ligand. From these two observations, we ascertain that the activity of the Ste2 complex, which is presumed to be higher in the presence of ligand, increases along with the interprotomeric distances of the quaternary structure. Therefore, the potential energy wells corresponding to the tighter conformation (i.e., shortest *r*_1_ and *r*_2_ distances) are drawn to the far left of the energy landscape diagram in Figure 5 (indicated by the solid blue arrows); the change in relative abundance of this inactive basal state conformation upon ligand binding is depicted in Figure 5a and b as a decrease in the depth of the energy well attributed to this inactive state. Likewise, we attribute the two longest *r*_1_ and *r*_2_ distances to those of the fully active conformation and draw the associated energy wells to the far right of the potential energy landscape of Figure 5.

We find not only that the interprotomeric distances of the Ste2 quaternary structure increase as the receptor switches from the basal state to the fully active state, but also that such increase does not scale equally in both dimensions. We see from a comparison of the *r*_1_ and *r*_2_ distances listed in Tables 1 and 2 that the change in distances between protomers occurring along one dimension of the parallelogram (*r*_1_ distance) is more pronounced than the change occurring along the other (*r*_2_ distance). If the adjustment of the position/orientation of the TM domains in individual Ste2 protomers is such that the specific TMs which serve as binding interfaces between protomers along *r*_1_ flare out more than those TMs involved in the binding interfaces between protomers separated by *r*_2_, then one could explain why elongation was more pronounced along one dimension of the tetramer versus the other (see Supplementary Figures S3 and S7).

The second significant observation, which may be made upon inspection of the *r*_1_ distance histograms, is that the Ste2 quaternary structure can populate multiple quaternary conformational substates. Specifically, the extremely broad peak seen in Figure3a (labeled Peak M) separated into two well defined narrow peaks when Ste2 was exposed to ligand (labeled Peak 2 and 3 in Figure 3c). Following our hypothesis that Peaks 1 and 4 correspond to inactive and fully active states, respectively, we attribute the peaks located between Peak 1 and 4 to semi-stable partially active conformational states. This broad peak (Peak M) is likely due to the protein complex shuttling between two partially active intermediate conformations which are separated by a low, flat energy barrier. Our interpretation of this situation is represented by the dashed black line in Figure 5a, which shows a low energy barrier between the two energy minima identified as states 2 and 3. The receptor complex can populate these conformational substates even in the absence of ligand, as is evident from the high relative abundance value calculated for Peak M in Table 1. However, because of the low energy barrier between these intermediate states, thermal perturbations would allow the quaternary structure to sample a multitude of configurations between the two states (i.e., Peaks 2 and 3) on a time scale which is faster than can be captured using our instrument. Only when ligand is added are these states “locked in” for long enough intervals to be captured via FRET spectrometry, an effect which both deepens the minimum and increases the height of the barrier between the two partially active states^50^. A similar phenomenon of partially active states being populated in the absence of ligand has been observed in the adrenergic 2A receptor, where four conformational states (two inactive and two active) were always present^32^. Agonist binding simply shifted the equilibrium of the conformations, with the higher activity states becoming more populated upon ligand binding.

One potential explanation as to why the Ste2 quaternary structure samples multiple intermediate states in the presence of ligand may be a result of sequential ligand binding (multiple step binding process), which has been shown to occur for other GPCRs^31^. Sequential ligand binding occurs when the contacts between the receptor and ligand do not form at the same time. Each contact which is formed between receptor and agonist results in a different conformational intermediate of the receptor, with the receptor becoming more active with each added contact. A recent study using atomic three-dimensional homology-based simulations showing Ste2 ligand binding occurs between 26 residues primarily within transmembrane helices H1, H5, and H6 which may suggest that the three states identified as 2, 3 and 4 in Figure 3c and Figure 5a might actually be a result of sequential binding of the agonist to one of each of these helices (in an order which is currently unknown^78^). Each new contact which is made between the ligand and a particular helix would result in a conformational change of the Ste2 receptor, and hence induce a change in the quaternary structure as well. Another GPCR, the β_2_ adrenoceptor, shows that upon each sequential step of agonist binding, some stabilizing intramolecular interactions are broken which allow for a higher probability that additional contacts may form as the receptor explores its conformational landscape^31^. The plasticity of intramolecular interactions (i.e., changes of distances between TMs, or forming new bonds/breaking previously formed bonds) as a GPCR binds at multiple steps with an agonist may provide further evidence of multiple activity levels within the tertiary structure, which is a concept we draw upon to describe similar effects at the quaternary level.

An alternative explanation for the existence of multiple quaternary substates may lie in the concept of membrane potential depolarization. Previous studies have shown that the muscarinic receptor type 2 (M2) displayed tertiary conformational changes which were induced by changes in membrane potential, particularly in the ligand-binding pocket, leading to a number of conformational states^79–80^. By similarity, the muscarinic receptor type 1 (M1) was assumed to bind agonist for a depolarized membrane which resulted in an acceleration of the transition from a quiescent to an active conformation, assuming multiple states along the path of activation^81^. We propose that by extension, the existence of multiple inactive and active states (including partially active substates) at the tertiary level may be reflected at the quaternary level as well.

One potential effect that may be ruled out as the cause of Ste2 existing in multiple conformational substates relates to the preassembly of G-proteins and their pre-coupling with GPCRs, namely the possibility that a different number of G-proteins are pre-bound to the GPCR^82^. While Li et. al. showed that preassembly of the heterotrimeric G-protein is highly independent of ligand presence, the Gα-subunit alone is found in low abundance and is unlikely to bind to receptors at a basal level. Their findings also suggest that a basal level of preassembled Gαβγ bound to receptors in the absence of ligand is much lower compared to when it is present. Li’s findings provide evidence that quaternary structure substates observed in this work exist independently from basal levels of G-proteins binding to the GPCR in various combinations regardless of whether the ligand is present or not.

## 5. CONCLUSIONS

While the findings presented in this work remain consistent with the previous finding that Ste2 forms parallelogram-shaped tetramers at relatively low concentrations within the plasma membrane, the more elaborate set of experiments, larger sample size, and improved methodology resulted in the unprecedented experimental detection of Ste2 quaternary structure substates in living cells. We also found that, when Ste2 was exposed to agonist ligand, the inter-protomeric distances increased, suggesting a quaternary structure conformation that corresponds to the receptors being in a more active state. We ascribed the different quaternary conformational substates to different levels of activity of the receptors, which is consistent with current knowledge on the conformational states populated by the tertiary structure of individual receptors.

The work presented here demonstrates the power of FRET spectrometry to detect shifts in the abundance of specific quaternary conformational states upon the binding of an agonist. This methodology may be used to determine how different ligands (such as agonists, inverse agonists, antagonists) modulate the conformational substates of the quaternary structure of any GPCR. Furthermore, by combining FRET spectrometry with the complete atomic crystal structure of the GPCR under study, information which is currently lacking for the Ste2 receptor, the binding interfaces and binding energies between protomers within the oligomer could be ascertained ^28^. Furthermore, availability of crystal structure information would facilitate identification of a more direct connection between the hinging motion of the receptor transmembrane domains and the receptor quaternary structure, which would allow probing the former by monitoring changes in the latter in the absence and presence of various ligands.

## ACKNOWLEDGMENTS

The optical micro-spectroscopy imaging facility used for this research was developed with support of the National Science Foundation, Major Research Instrumentation Program (Grant No. PHY-1126386 awarded to V.R.). This work was partially funded by secondary support of the National Science Foundation, Major Research Instrumentation Program (Grant No. PHY-1626450 awarded to V.R.) and by the National Science Foundation, Physics of Living Systems Program (Grant No. PHY-1058470 awarded to V.R.). We thank Ionel Popa for access to his protein purification facility and Claudiu Gradinaru for stimulating discussions.

## SUPPORTING INFORMATION

Supplementary methods and data may be found online at _____.

## Footnotes

1 In both the absence and presence of ligand, the best fit for models consisting of three peaks and four peaks had reduced *χ*^2^ of 1.61 and 0.15, respectively.

2 In both the absence and presence of ligand, the best fit model had a reduced *χ*^2^ of 0.70 and 1.05, respectively, when simultaneously fit using equivalent mean positions of the two individual Gaussian curves (labeled Peaks).

## Notes

### Competing Interest Statement

The authors have declared no competing interest.

